# How to design 1000-plex mass tags using the differential mass defect

**DOI:** 10.1101/2025.10.22.679607

**Authors:** Harrison Specht, Michael P. Agius, Kevin McDonnell

**Author notes:** These authors contributed equally (Michael P. Agius, Kevin McDonnell). Data and code: parallelsq.org/tagdesign.

## Abstract

Multiplexing samples in mass spectrometry-based proteomics has long been accomplished by isotopologues of small molecules. These chemically-identical “mass tags” conjugate to peptides to encode samples with different mass offsets for parallel analysis. The current state-of-the-art for multiplexing with non-isobaric mass tags was recently improved from 3-plex to 9-plex, but what is the largest plex size that can be reasonably achieved with current technology? A full answer to this question requires evaluating current mass spectrometry hardware, facets of which have been well-investigated by others. However, it may be underappreciated that multiplexing 1000s of samples with mass tags does not actually require 1000s of isotopes, or 1000s of synthesis steps to create. Non-intuitively, high plex mass tags can require relatively few different isotopes. The focus of this exposition is to characterize the potential of the *differential mass defect* to create tens to over a thousand isotopologues of small molecules and how careful combinations of these small molecules can combinatorially scale the plex size to minimize synthetic steps. Importantly, we show that plex sizes in the hundreds, an order of magnitude greater than state-of-the-art, are achievable using molecules comparable in size to existing commercial tags, and that going beyond hundreds may require larger molecules. Approaches to achieve high-plex proteomics will almost certainly require using the differential mass defect, so we hope this exposition serves to accelerate progress in reagent development to achieve high plex proteomics.

## Introduction

Global proteome characterization by mass spectrometry is rapidly becoming common practice across biological and drug discovery sectors^1^. However, sample throughput still remains a significant bottleneck for expanding into translational and population based studies^2–4^. Multiplexing samples was popularized by the success of isobaric mass tags, including TMTpro, which enables 35 samples to be analyzed in a single run with data-dependent acquisition (DDA) methods^5–8^. However, with depth of coverage significantly enhanced using data-independent acquisition (DIA) methods, combining this with mass tag sample multiplexing has the potential to give even greater increases in throughput^9^. DIA and non-isobaric mass tags (i.e. mTRAQ) were shown to allow 3-sample multiplexing capabilities and were coined “plexDIA”^9^. Recently, we reported a 9-plex mass tag for plexDIA, PSMtag^10^. PSMtag reaches larger plex sizes from an increase in dopable sites reflected in its larger molecular weight (327 Da). However, increasing plex size simply by adding more dopable sites will certainly reach a practical limit, so we sought to understand if high plexes could be created with limited dopable sites for application in either DDA or DIA workflows. We investigated and present here how the *differential mass defect* can be utilized to create high plex mass tags with limited dopable sites. Importantly, we will not include in this exposition simulations for the depth of proteome coverage for different mass defects and mass spectrometer resolutions, as this has been described by others previously^11^. We will also not examine the effects of increased ion flux associated with multiplexing, in the mass and time domains^12^, and associated with fast gradients analyzing hundreds of samples-per-day (SPD)^13^. We hope this exposition in exploiting the differential mass defect to creating high plex reagents will accelerate progress in reagent development.

### The “mass defect” and why we found a new term helpful

The term “mass defect” has a long history with various fields using it to describe slightly different concepts, leading to some confusion being noted^14,15^. In physics, the “***nuclear* mass defect**” is defined as the difference between the empirical mass of a molecule and the sum of the masses of its constituent parts: neutrons, protons, and electrons. Its precise value can be calculated using Einstein’s equation *E* = *mc*^216^. In mass spectrometry, the “***chemical* mass defect**” has adopted a different meaning, being used to describe the difference between the observed mass of a molecule and its nominal mass^17,18^. Here, the **nominal mass** is the sum of the integer masses of the constituent elements (number of nucleons). With mass spectrometers being one of the first ways to observe the difference between a molecule’s actual mass and nominal mass, this metric has a long history in the field. In 1927, Aston described the “packing fraction” as the ratio of an atom’s chemical mass defect to its nominal mass^19^. More recently, further variations on this metric have also been introduced, such as the Kendrick mass defect^20^. The Kendrick mass defect has been useful in highlighting important patterns within the data^21^, and we hope that the metric defined in this exposition will also prove useful. Within their respective fields, the nuclear mass defect and the chemical mass defect are both typically referred to as just the “mass defect”. Although they are ultimately the result of the nuclear binding energy, acknowledging the difference between them is important. The nuclear mass defect is an absolute parameter, while the chemical mass defect is a relative value dependent on the scale used (typically ^12^C). Using more specific terms, as we have done here, will avoid confusion between the two meanings. A more comprehensive comparison of the different uses of mass defect is performed by Pourshahian^14^. In this exposition, from now on any use of “mass defect” alone will refer to the nuclear mass defect.

### Defining the “differential mass defect”

An example of the mass defect is shown in Fig. 1a, where for a ^12^C atom the difference in mass is approximately 10 mDa. This defect is also distinct for different elements and their isotopes. Indeed, this discrepancy has previously been incorporated into mass tag design for multiplexing proteomic samples. By combining mass defects from different elements, one can create tag isotopologues with sub-dalton mass differences^22,23^. In our own studies designing mass tags for multiplexing proteomic samples^24^, we found a direct measure of this phenomenon to be helpful, a metric we term the “differential mass defect”. The **differential mass defect** is defined as the difference in mass between molecules of the same chemical structure (isotopologues) comprising the same number of protons, electrons, and neutrons, but different elemental isotopes. Although these isotopologues share the same number of neutrons, some are located in different elements, resulting in a different molecular mass. The change in the location of the neutrons modifies the summed binding energy of the different isotopologues. Examples of the differential mass defect of various isotopologues are shown in Fig. 1a. Unlike other variations of the mass defect terminology, this metric directly describes the difference in masses of the mass tags that will be used to differentiate multiplexed samples in the mass domain.

**Figure 1.**
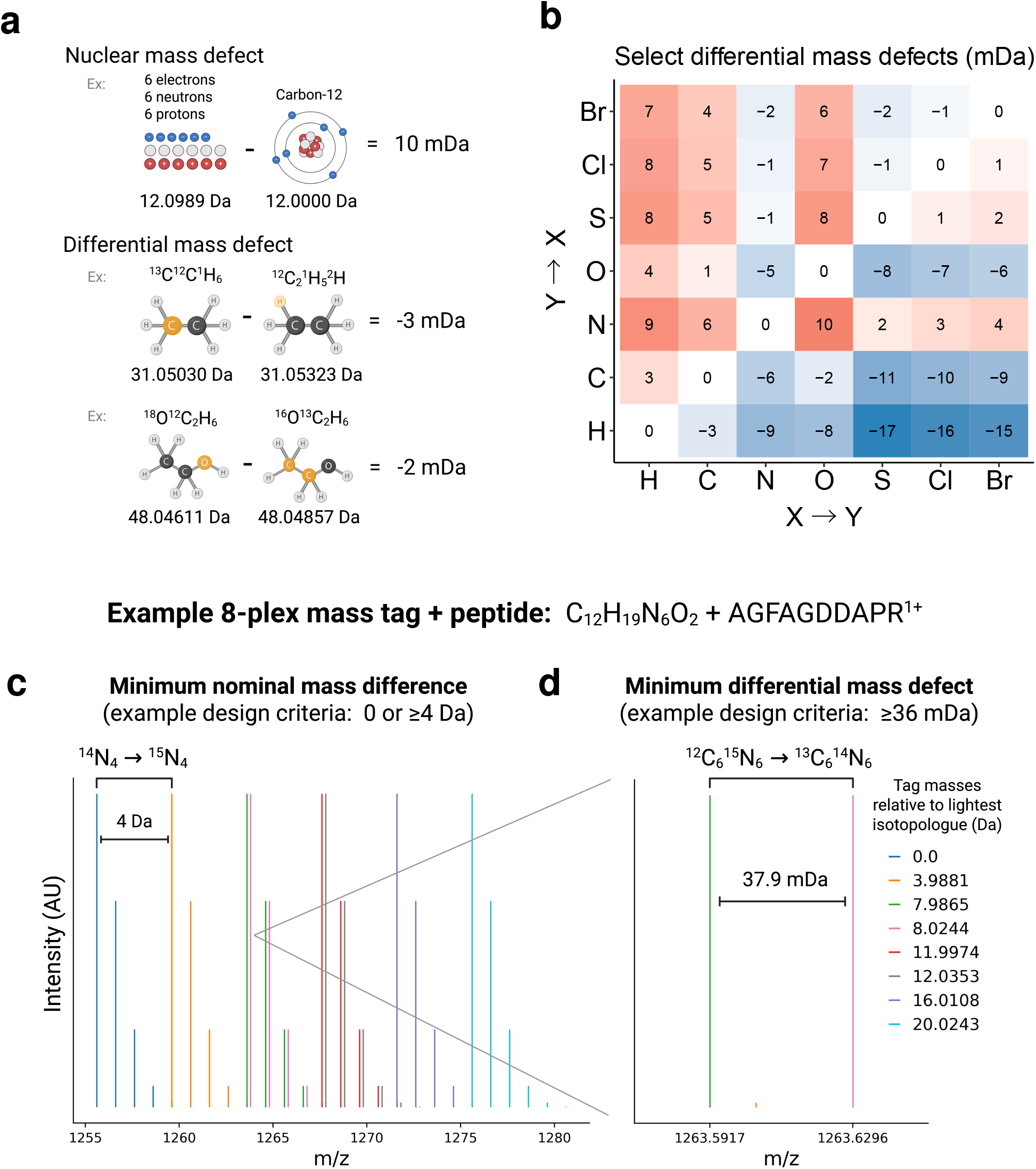
Describing the differential mass defect. **a**, The nuclear mass defect: the difference in mass between the whole atom and its constituent parts. The differential mass defect: the difference in mass between molecules comprising the same number of protons and neutrons and chemical structure, but different elemental isotopes. **b**, Selected differential mass defects created between combinations of 1-2 elements. Each difference is created using the most common isotopes of each element, for example ^37^Cl. **c**, The different isotopic distributions for an 8-plex tag with a base composition of C_12_H_19_N_6_O_2_ attached to the peptide AGFAGDDAPR. The nominal mass difference of 4 Da between the monoisotopic peaks of the first and second channels is obtained by replacing ^13^N_4_ with ^14^N_4_. mDa differences are exaggerated for visibility. **d**, The monoisotopic peaks for the third and fourth channels for the same tag and peptide are highlighted. A differential mass defect of 37.9 mDa is obtained by replacing ^12^C_6_^15^N_6_ with ^13^C_6_^14^N_6_.

## Results

### Creating differential mass defects from a variety of elements

Multiplexing in bottom-up mass spectrometry has typically used nominal mass nomenclature to define each channel e.g. Δ0, Δ4 and Δ8 signifying the nominal mass difference in Daltons from the base tag. However, when considering the differential mass defect, one can have multiple channels at the same nominal mass. Isotopologues with the same nominal mass, i.e. those comprising the same number of protons, electrons and neutrons, can be created from different combinations of elemental isotopes. For example, ^18^O^12^C_2_^1^H_6_, ^16^O^13^C_2_^1^H_6_, and ^16^O^12^C_2_^2^H_2_^1^H_4_ all have the same nominal masses. However, these isotopologues have different actual masses due to the distinct binding energies of the different atoms which hold additional neutron(s). To more systematically view this phenomenon, differential mass defects between pairwise combinations of select elements were calculated and depicted in Fig. 1b. For example, in a compound where one ^37^Cl and two ^1^H have been exchanged for one ^35^Cl and two ^2^H, the differential mass defect is -16 mDa. However, this does not represent the limit for the differential mass defect of a compound. Should the compound contain 2 such opportunities for exchange (a total of two Cl for four H), the differential mass defect would be (2)*(-16) = -32 mDa. Similarly, exchanging 6 carbon and nitrogen isotopes allows creating a total of (6)*(6 mDa) = 36 mDa. In summary, the differential mass defects depicted in Fig. 1b are able to be combined to create larger (or smaller) differential mass defects. This analysis was limited to select stable elemental isotopes that are widely-accessible: ^12^C, ^13^C, ^1^H, ^2^H, ^16^O, ^18^O, ^32^S, ^34^S, ^35^Cl, ^37^Cl, ^79^Br, and ^81^Br. The differential mass defects are symmetrical around the diagonal when 1 or 2 neutrons in *1 atom* are exchanged for the same number of neutrons in *1 other atom*. The differential mass defects are not symmetrical around the diagonal when 2 neutrons in *1 atom* are exchanged for the same number of neutrons in *2 other atoms*. This is due to the difference in the binding energies of adding 2 neutrons to 1 nucleus compared to 2 neutrons separately to 2 nuclei.

### Small molecules can create 10-1000 plex mass tags using the differential mass defect

To explore how the differential mass defect expands the plexing capabilities of mass tags, we calculated the masses, and therefore possible plex sizes, of all possible isotopologues for compounds resembling current mass tags. We tested a range of empirical formulas following Lipinski’s rule (a reasonable starting point for synthetic small molecules) with the addition of bromine and chlorine, and the lightest isotopologue being less than 600 Da. A molecular weight of 600 Da is slightly less than the mass added by a current commercial tag (TMTpro^25^ to lysine-containing peptides), and served as a reasonable cutoff. Two additional criteria that strongly inform the plex size achievable by a given compound include: 1) the **minimum difference in nominal mass** desired and 2) the **minimum differential mass defect** resolvable. Successful commercial mass tags have imposed a minimum nominal mass difference of 4 (Da) between reagent masses (Fig. 1c). This induces minimal overlapping signal from one plex channel into adjacent plex channels caused by each and every peptide’s naturally-occurring isotopic envelope. Recent advances in deconvolving peptide envelopes have demonstrated that a minimum nominal mass difference of 2 (Da) also allows accurate identification and quantification of proteomes^26^.

In parallel to the channels separated by the nominal mass differences, one can introduce alternative series of channels separated by the differential mass defect (Fig. 1d). The minimum differential mass defect that is practical for deep proteome coverage and accurate quantification is a function of mass spectrometer resolution (speed) and protein dynamic range. Some trade-offs necessary to use less than 36 mDa minimum differential mass defect are characterized by Merrill *et al*^11^. While popular commercial non-isobaric mass tags have not used the differential mass defect, an 18-plex using 36 mDa resolution has been demonstrated though it required introducing some labels in cell culture^22,27^. Resolution of differential mass defects of 12 mDa and 6 mDa across multiplexed proteomes represents a technical challenge, but offer significant benefits, with many candidate mass tags allowing 100s of samples to be plexed as shown in Fig. 2a.

**Figure 2.**
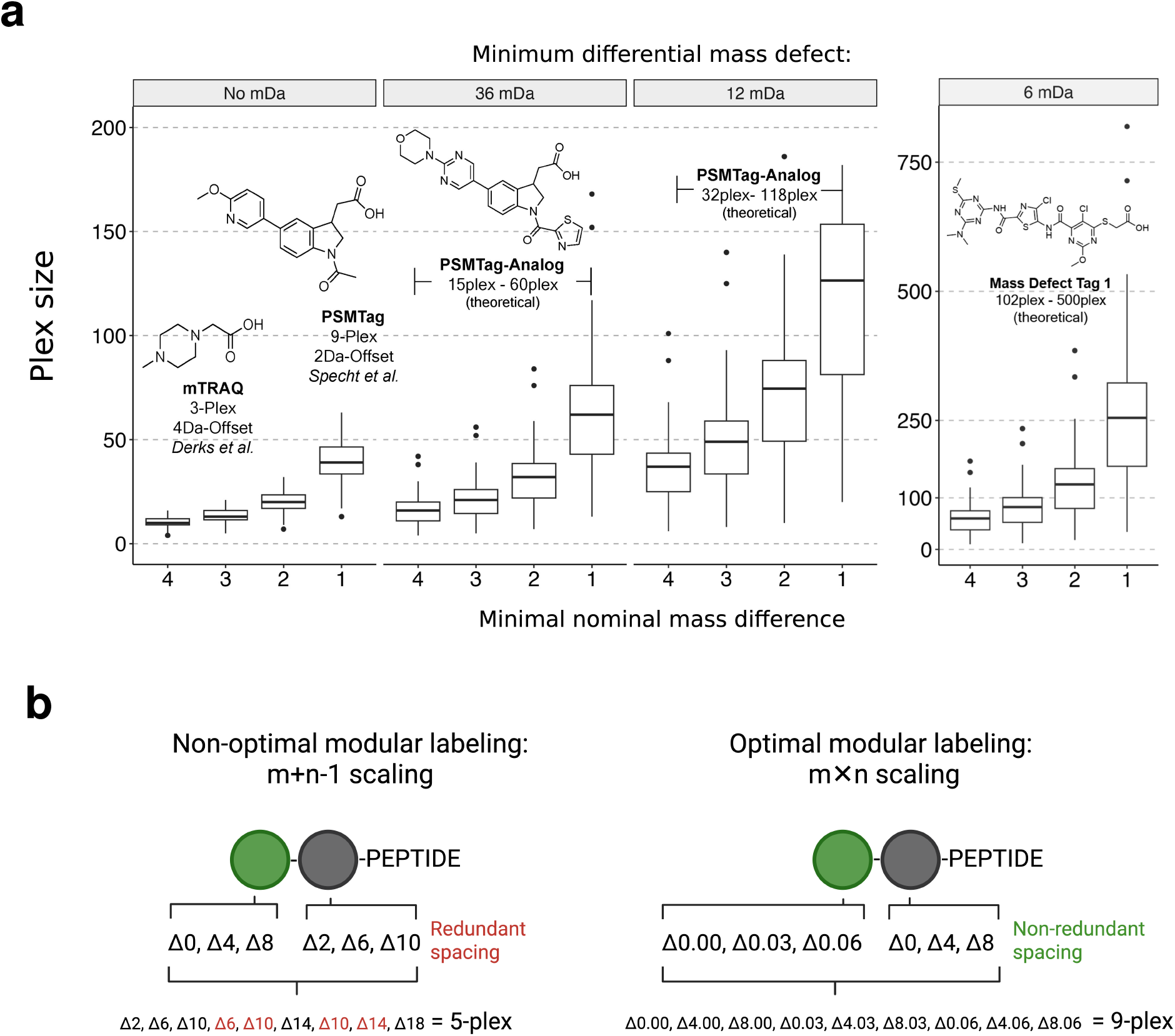
Designing mass tags to efficiently use the differential mass defect. **a**, Calculating the theoretical plex size for molecules *<*600 Da molecular weight and molecular formula generally following Lipinski’s rule as a reasonable starting point to represent synthesizable space: C_5-50_H_10-100_O_0-10_N_0-10_S_0-5_Br_0-2_Cl_0-2_. Numbers of elements per molecule were randomly sampled with uniform probability over the stated ranges, except for the numbers of Br and Cl, which were both sampled from the set { 0,0,0,0,0,0,0,0,1,2 }to have a realistic number of compounds containing either. **b**, Non-optimal and optimal combinations of masses in modular mass tag reagent design. Optimal combinations with *m* × *n* scaling can be achieved when the set of differences between the *m* versions of module 1 are disjoint to the set of differences between the *n* versions of module 2.

We offer as an example a variation of PSMtag (PSMtag-Analog) encoding more isotopes while additionally leveraging the differential mass defect. These modifications to the PSMtag scaffold achieve a 15 to 60-plex across the range of minimum nominal mass differences of 1 to 4 while using a minimum differential mass defect of 36 mDa. This higher plex mass tag maintains a reasonable molecular weight while utilizing commercial building blocks with potential for isotope enrichment. Additionally, we propose a novel mass tag (Mass Defect Tag 1) that offers *>*500-plex size when using 1 Da minimum nominal mass difference and 6 mDa minimum differential mass defect. Synthesis of the lightest isotopologue of Mass Defect Tag 1 is reasonable using commercially available building blocks, however full isotopic enrichment may rely on further methodology development. To push this strategy to its limits, we loosened up our molecular weight and elemental restrictions. While still *<*1000 Da and using commercially available building blocks, we are able to achieve *>*1000-plex with a tag of elemental formula C_33_H_44_N_15_O_5_S_3_Cl_3_ with a minimum nominal mass difference of 1 Da and minimum differential mass defect of 6 mDa. These reagents described in Fig. 2a represent proof-of-concept mass tags where the differential mass defect can unlock significantly larger plex sizes. Additionally, this analysis makes it clear that going beyond plex sizes in the hundreds will require significantly larger molecules.

### Modularization can allow combinatorial scaling of plex size in fewer synthetic steps

Synthesis of 100s to 1000s of mass tag isotopologues need not require 100s to 1000s of synthetic schemes if synthesis of the mass tags can be modularized. Fig. 2b shows a hypothetical mass tag with two subunits. The two subunits can be combined prior to peptide conjugation or after one of the subunits is conjugated to the peptide, the resulting masses, and thus plex size, is the same. However, not all modularization schemes result in fewer synthesis schemes than plexes, as shown in the “non-optimal” example in Fig. 2b. In this example, each of the two subunits have 3 possible mass configurations. If the subunits are chemically distinct and thus require different synthetic pathways to dope, at least 6 schema (3+3) are needed but only achieve a 5-plex due to redundancy of the mass offsets. This equally applies to dalton or subdalton mass offsets. However, by introducing different offsets in the subunits, redundancy of the combined masses can be avoided. For subunits with *m* and *n* respective mass offsets, *m* × *n* scaling of the plex size to modular subunits (*m* + *n*) can be achieved, as shown in the “optimal” example in Fig. 2b. The set of mass differences between the configurations *within* each subunit (0.03 Da, 0.06 Da and 4 Da, 8 Da in Fig. 2b) must be different between each subunit to allow all combinations to produce non-redundant masses. This design criteria achieves the largest plex size with the minimum number of modular subunits. In practice there are numerous chemical building blocks that can offer modularity to enhance plex size through isotope mass defects. As carbon and nitrogen isotopes are commonly commercially available at high isotopic purities, reagents like triazines, triazoles, and guanidine offer carbon to nitrogen ratios suitable for differential mass defects exploration. As a modular approach, these functional groups can be added during isotopologue synthesis or even applied ‘on-peptide’, post an initial labeling step, to create the differential mass defect (Fig. 2b). There are numerous bioorthogonal reactions to explore tag design including click-chemistry, oxime formation, and sulfur michael-additions. Such modular steps can be implemented either during isotopologue synthesis, or step-wise through bioorthangoal reactions, after first functionalization of bulk peptides (e.g. amine functionalization via NHS-ester chemistry).

## Discussion

### Notable caveats in simulating mass tag plex size

The exploration of mass tag plex sizes for a range of hypothetical molecules in Fig. 2a has several notable caveats. 1) It assumes all sites are available for isotope incorporation (except for all hydrogen sites, which were omitted completely due to the possibility of inducing retention time shifts, and the carboxy -OH). High isotopic doping percentage is not without precedent: TMTpro dopes 100% percent of carbons and nitrogens outside the reactive portion of the molecule^28^. Still, this exploration should be taken as representing an optimistic case for any of the given molecules. We find more value in interpreting the trends than the precise values for this plot. 2) The exploration also assumes that synthesis of the mass tag isotopologue without any higher mass isotopes is desirable. However, the largest set may or may not use the lightest isotopologue. The question of determining which set of isotopologues allows the largest plex size is mappable to the problem of finding the largest independent vertex set in a graph, where the graph is constructed by connecting mass tag isotopologues that are less than the minimum nominal mass difference or less than the minimum differential mass defect difference. An implementation of solving this problem for any given molecular formula is offered in the code base for this study, but is computationally expensive.

### Conclusion

In summary, we present the “differential mass defect” as a useful metric to consider for mass tag design, we offer examples of differential mass defects of common element pairs, we explore the range of plex sizes achievable by small molecules with and without employing the differential mass defect, and we propose modular mass tag design principles that allow for more efficient reagent production. We show that plex sizes in the hundreds, an order of magnitude greater than state-of-the-art, are achievable using molecules comparable in size to existing commercial tags, and that going beyond plex sizes in the hundreds may require larger molecules.

## Acknowledgments

We thank Nikolai Slavov and Maddy Yeh for feedback on this article. PTI is a Convergent Research Focused Research Organization (FRO) and has received support from Eric and Wendy Schmidt as well as Griffin Catalyst.

## Code availability

The code used for data analysis and figures is at: github.com/ParallelSquared/tagdesign

## Writing

All authors approved the final manuscript.

## Methods

All analysis and figures are reproducible from scripts written in the R coding language contained in the repository: github.com/ParallelSquared/tagdesign

### Calculating select differential mass defects

Elemental masses were derived from the Ame2012 atomic mass evaluation^29^. The stable elemental isotopes used in the calculations were: ^12^C, ^13^C, ^1^H, ^2^H, ^16^O, ^18^O, ^32^S, ^34^S, ^35^Cl, ^37^Cl, ^79^Br, and ^81^Br. The approach allows for 1 or 2 neutrons in *1 atom* to be exchanged for the same number of neutrons in *1 other atoms* or for 2 neutrons in *1 atom* to be exchanged for the same number of neutrons in *2 other atoms*. All such combinations of the above elements and their isotopes were considered. The calculations are performed in *select mass defects*.*R*.

### Estimating plex size from molecular formulas

The calculation of the maximum plex-size achievable for a tag molecule can be modeled as finding the maximum independent set of vertices in a graph. For a given tag molecular formula, one can generate the graph *G* = (*V, E*) where each vertex *v* is an isotopologue of the specified tag with a mass *m*_*v*_. An edge *e* is present between vertices *v* and *w* when the mass difference between them is less than the minimum mass defect (MMD) allowed:

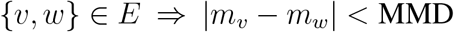

*or* the difference between *m*_*w*_ and any of the carbon isotopes of *v* less than the minimum nominal mass difference (MNM) from *m*_*w*_ are also less than minimum mass defect. This can be written as follows:

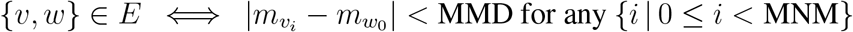

where 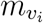 is the mass of the *i*^*th*^ carbon isotope of *v*, 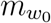 is the base mass of *w*, and *m*_*v*_ *< m*_*w*_.Carbon isotopes are used to model the expected natural isotopic patterns of the peptides the tags will be attached to.

For *G*, a maximal independent set *S* is defined as a set of vertices where for each vertex only one of the following conditions is met:

1. *v* ∈ *S*
2. *N* (*v*) ∩ *S*≠ ∅ where *N* (*v*) is the neighbors of *v*.

The maximal independent set with the largest number of vertices denotes the maximum independent set of *G* and is the maximum plex-size for that molecular formula. The ‘igraph’ package^30–32^ was used to find these analytical solution(s) as shown in *example ivs solver*.*R*, but this approach is computationally expensive.

A more computationally inexpensive algorithm for estimating plex size from molecular formulas proceeds as follows: first by enumerating all possible isotopic variants of a given molecular formula; then computing their exact masses; then filtering the list of all possible exact masses by criteria for minimum differential mass defect (MMD) and minimum nominal mass defect (MNM), starting with and keeping the lightest isotopologue. This approach is documented in *plex sizes*.*R* and was used in the creation of Fig. 2a.

### Example 8-plex mass tag + peptide

The isotopic envelope for peptide precursor ion AGFAGDDAPR (1+) was simulated using the ‘OrgMassSpecR’ package for R. The masses of the peptide isotopologues were then added to the masses of the mass tag isotopologues, calculated using the computationally inexpensive approach described above. The calculations are performed in *simulate single peptide envelope*.*R*.

